# Abated immune responses against recombinant hepatitis-b vaccine by chitin in mice

**DOI:** 10.1101/2021.03.25.436808

**Authors:** Parker Elijah Joshua, Nnaemeka Emmanuel Ayogu, Timothy Prince Chidike Ezeorba, Damian Chukwu Odimegwu, Rita Onyekachukwu Asomadu, Uzochukwu Gospel Ukachukwu

## Abstract

Majority of recombinant protein vaccines employ alum-based adjuvants to boost their immunogenicity in subjects. However, the safe-profiled alum-based adjuvants demonstrate relative ineffectiveness with these recombinant vaccines. This work investigated the adjuvant potential of chitin with recombinant Hepatitis-B protein vaccine (HBsAg) and its possible toxicological effects. Six to eight weeks old female albino mice, which were randomly and equally distributed into five groups were used for this study. Blood collection was done before each vaccination schedule and the samples were used for analysis. Results revealed that IgG and IgG_1_ titres of mice administered 2 doses of HBsAg-Chitin formulation was significantly (p < 0.05) lower than those administered 3 doses of HBsAg and was not significantly (p > 0.05) higher than those administered 2 doses of HBsAg. However, a progressive increase in the anti-HBsAg titres in mice that received the HBsAg-Chitin formulation was observed as days came by. Activities of alanine and aspartate aminotransferases of all experimental groups, including their liver weights, inferred that the HBsAg-Chitin formulation exert no toxic effect on the liver. The findings of this research revealed that chitin demonstrated a relatively unimpressive adjuvant activity with the recombinant hepatitis-B protein vaccine.

**GRAPHICAL ABSTRACT:** 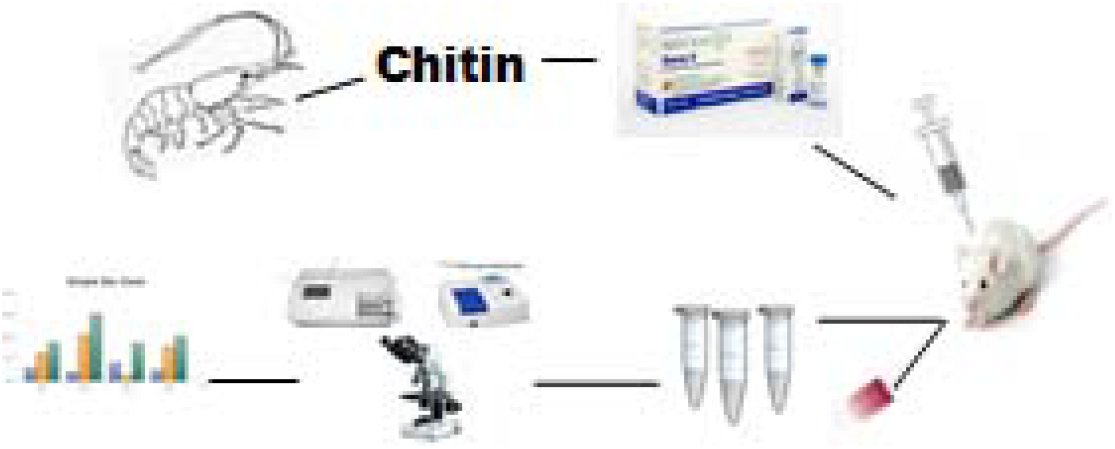

## 1. Introduction

Hepatitis-B virus infection is one of the chief causes of liver cancer and cirrhosis development and vaccines have demonstrated to be potential interventions for the prophylaxis of such infectious disease^1)^. According to O’Hagan and Gregorio (2009), contemporary vaccines in development contain highly purified antigens or recombinant proteins, representing pathogen subunits^2)^. Highly purified vaccines are usually administered to subjects in combination with adjuvants to enhance their immunogenic potentials^3)^. An immunologic adjuvant is any material that could quicken, protract, or improve antigen-specific immune responses following its combination with antigens^4)^; however, it is not expected to elicit an immune response in isolation. Studies have shown that vaccine-adjuvant formulations are promising and have remarkable potentials to stimulate immune responses that mobilize and activate dendritic cells (DCs), macrophages, and lymphocytes to eliminate pathogen of interest^5)^. Highlights of the outcome of application of adjuvants in vaccine development include; improving functional and total antibody titre, induction of potent cellular immunity, enhancing effective immune responses in infants and aged populations, reduction of the number of booster doses and the effective dose of the vaccine antigen, enhancing mucosal immunity and widespread redemptive immune responses^4)^. Recombinant Hepatitis B vaccine contains aluminum-based adjuvant that helps to boost its immunogenic profile, but has been exhibited weakness in activating the T-helper 1 mediated immune response responsible for elimination of viral infections intracellularly^5)^. Therefore, several vaccine booster doses are necessary to attain optimum immunity, even though the response is non-lasting^6)^. Furthermore, alum adjuvant-based vaccines have been shown to be unsuccessful because it elicits primarily T-helper 2 mediated response which is ineffective for vaccination against many types of intracellular pathogens^3)^. Chitin, a long-chain polymer of a derivative of glucose, N-acetylglucosamine has found widespread application in medicine and pharmacy as a drug-delivery system due to its biocompatibility, ability to adhere to mucosal layers, non-toxicity and biodegradability^7)^. According to Li *et al*. (2013), being found generally in lower organisms, Chitin was said to be the second most abundant natural polysaccharide^7)^. Previous studies reported that chitin and some of its derivatives may possess antiviral, antimicrobial and adjuvant activity^8,9)^. The potential to mobilize and stimulate the effectors of the innate and adaptive immune systems by chitin has been associated with variations in its complexity and size^10)^. Intravenous administration of chitin particles induced a significant priming effect in natural killer (NK) cells and alveolar macrophages in mice^11)^. Again, phagocytosable small-sized chitin particles were shown to induce the expression of cytokines such as interleukin-12 (IL-12), tumor necrosis factor-a (TNFa), and IL-18 by alveolar macrophages, resulting in predominant production of interferon-γ (INF-Y) by natural killer cells^12)^. Furthermore, production of cytokines (IL-12) by the alveolar macrophages was shown to be activated via mannose receptor-mediated phagocytosis^12)^. The aim of this study is to investigate adjuvant activity of chitin when constituted with a recombinant Hepatitis-B protein vaccine (HBsAg) and also the toxicological effect this formulation would exact on the liver.

## 2.0 MATERIALS AND METHODS

### 2.1. Reagents/Chemicals

In this study, all reagents and chemicals used were of analytical grade and are listed as follows; 96% Acetone (JHD), Aspartate aminotransferase (AST) Kit (DiaLab, Austria), Alanine aminotransferase (ALT) Kit (DiaLab, Austria), Carbonate-Bicarbonate Buffer (pH 9.5), Phosphate Buffered Saline (pH 7.4), Fat-free Milk powder (Dano-Slim, Nigeria), Citrate Phosphate Buffer (pH 5.0), DMSO (JHD, China), methanol, Hydrogen Peroxide (JHD, China), 2 M H2SO4 (JHD), Tetramethylbenzidine (TMB) Tablets – (SIGMA, USA), Leishman stain, Tween 20 (SIGMA, USA).

### 2.2. Prawn

Frozen prawns, *Litopenaeus vannamei*, were purchased from Bonny beach market, Bonny Island, Rivers State, Nigeria and were submitted to the Department of Zoology and Environmental Biology, University of Nigeria for identification and validation.

### 2.3. Chitin Extraction

Chitin was extracted from the prawns using the method described by Islam et al. (2016) ^13)^. The fresh frozen prawn shells were washed with hot water (95°C), dessicated and then pulverized. Following pulverization, the shell powder was deproteinated in 5% NaOH solution, and demineralised with 1% HCl (w/v 1:10) to obtain chitin.

### 2.4. Preparation of the Adjuvant (Chitin Dispersion)

Chitin powder extract was dispersed in normal saline; 6mg of the powder was dispersed in 1ml of normal saline and vortexed for adequate hydration for 20mins.

### 2.5. Secondary Antibodies and Vaccine

The secondary antibodies used for the study were the goat anti-mouse IgG1, IgG and IgG_2a_ immunoglobulins (Southern Biotechnology, USA) conjugated with horseradish peroxidase. Euvax-B (LG chemical, South Korea), a recombinant hepatitis B protein vaccine (HBsAg) was employed as the vaccine candidate in this study.

### 2.6. Animal Procurement

Six to eight weeks old female albino mice, *Mus musculus*, were procured from the animal facility of the faculty of Veterinary Medicine, University of Nigeria and were acclimatized for 7 days under standard housing conditions with a 12 hour light/dark cycle. They received standard pellets and tap water *ad libitum*. The University of Nigeria regulations for laboratory animal use and the NIH guidelines for laboratory animal care and use were implemented while handling the laboratory animals as outlined by National Research Council, 1996^14)^.

### 2.7. Experimental/Study Design

The mice were divided into five (5) groups with five (5) animals per group and treatment was done as follows:

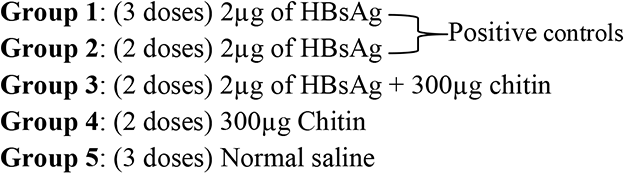

### 2.8. Immunization Protocol

The mice were vaccinated via the intramuscular route in a prime-boost fashion with 2μg of HBsAg (3 and 2 doses respectively), 2μg of HBsAg + chitin (300μg), chitin (300μg) only or normal saline (100 μL). Animals receiving 2 doses were vaccinated on days 1 and 15 and those that received 3 doses were administered on days 1, 15 and 22. Blood sample collection from the mice was done via retro-orbital plexus on days 0, 14, 21 and 28 and the sera obtained after centrifugation at 5000 rpm for 10 minutes. Equal aliquots of sera were frozen until further analysis.

### 2.9. Determination of Antibody Titres against Vaccine/Formulation

Baseline anti-HBsAg titre was determined by analysis of blood samples collected on day 0 and subsequently, antibody titre analysis of blood samples collected on days 14, 21 and 28 were conducted. Analysis of specific total antibodies (IgG), IgG1 and IgG_2a_ was done using an indirect ELISA method described by Ternette *et al*., (2007) with slight modification^15)^. 100μL of 20μg/mL r-HBsAg (from vaccine) in carbonate-bicarbonate buffer of pH 9.5 (coating buffer), was coated on ninety six (96-well) ELISA plates (BRAND plates®, Germany) and subsequently incubated overnight at 4°C. Washing of the wells was done thrice with 300 μL of washing buffer (PBS of pH 7.4 + 0.05% Tween 20) and in turn was blocked with blocking buffer (PBS of pH 7.4 + 5% Fat-free milk) at 37°C for 1 hour. 1:500 dilutions of sera (containing the primary antibodies) were prepared in antibody buffer (2% Fat-free milk in PBS-Tween). Following washing of the wells, 100μL of diluted primary antibodies was added to each well and incubated for 1 hour 30 minutes at 37°C. Another washing was done, and 100μL of 1:1000 dilution of secondary antibodies was added to each well and incubation was done for another 1 hour 30 minutes at 37°C. Following repeated washing, 100μL of substrate solution (TMB + hydrogen peroxide + citrate phosphate buffer, pH 5.0) was put into each well and the plates were placed in a dark enclave at room temperature for 20 minutes for sufficient colour development. A stop solution (50 μL of 2M H_2_SO_4_) was added after colour development. Following reaction stoppage, the optical density (OD) was determined at 450nm using an automated ELISA plate reader (GMI, USA).

### 2.10. Differential White Blood Cell Count

On days 0, 14, 21 and 28, the white blood cell differential count was analyzed according to the method described by Rodak and Carr (2016)^16)^. The differential white blood cell counts were expressed as a percentage.

### 2.11. Liver Function Test

Serum AST and ALT concentrations were measured using diagnostic kits (DiaLab, Austria). Assaying for AST and ALT activities was according to the methods described by Thomas (1998) and, Moss and Henderson (1999) respectively^17,18)^.

### 2.12. Determination of Body Weights

The mice were weighed throughout the duration of the study, from days 0 to 28, using an electronic weighing scale (HX502T) and their body weights were recorded appropriately.

### 2.13. Determination of Liver Weights

On the last day of the study (day 28), the mice were sacrificed and their livers were harvested and weighed using an electronic weighing scale (HX502T). The liver weights were recorded.

### 2.14. Statistical Analysis

Results were expressed as mean ± standard deviation [S.D]. Data were analyzed using one-way ANOVA and subjected to Duncan tests on SPSS (Statistical Product and Service Solution) vs.20 software. Differences between means of the treated groups and control were considered significant at p < 0.05. All figures were made using GraphPad Prism version 7.0.

## 3.0 RESULTS

### 3.1. Effect of HBsAg-Chitin formulation on IgG Titre of Mice

**Figure 1** showed that mice administered with 2 doses of HBsAg-Chitin formulation elicited a significantly (p < 0.05) lower IgG titre level when compared to mice administered with 3 doses of HBsAg (group 1) at days 14 and 21. However, mice administered with 2 doses of HBsAg-Chitin formulation exhibited a significantly (p < 0.05) higher IgG titre when compared to mice administered with 2 doses of HBV alone (group 2) on day 21. Interestingly, IgG titre increased progressively from days 0 to 21, in mice administered HBsAg-Chitin formulation. Moreover, IgG titres of mice administered 2 doses of HBsAg were not statistically significant from those that received 2 doses of HBsAg-Chitin formulation.

**Figure 1:**
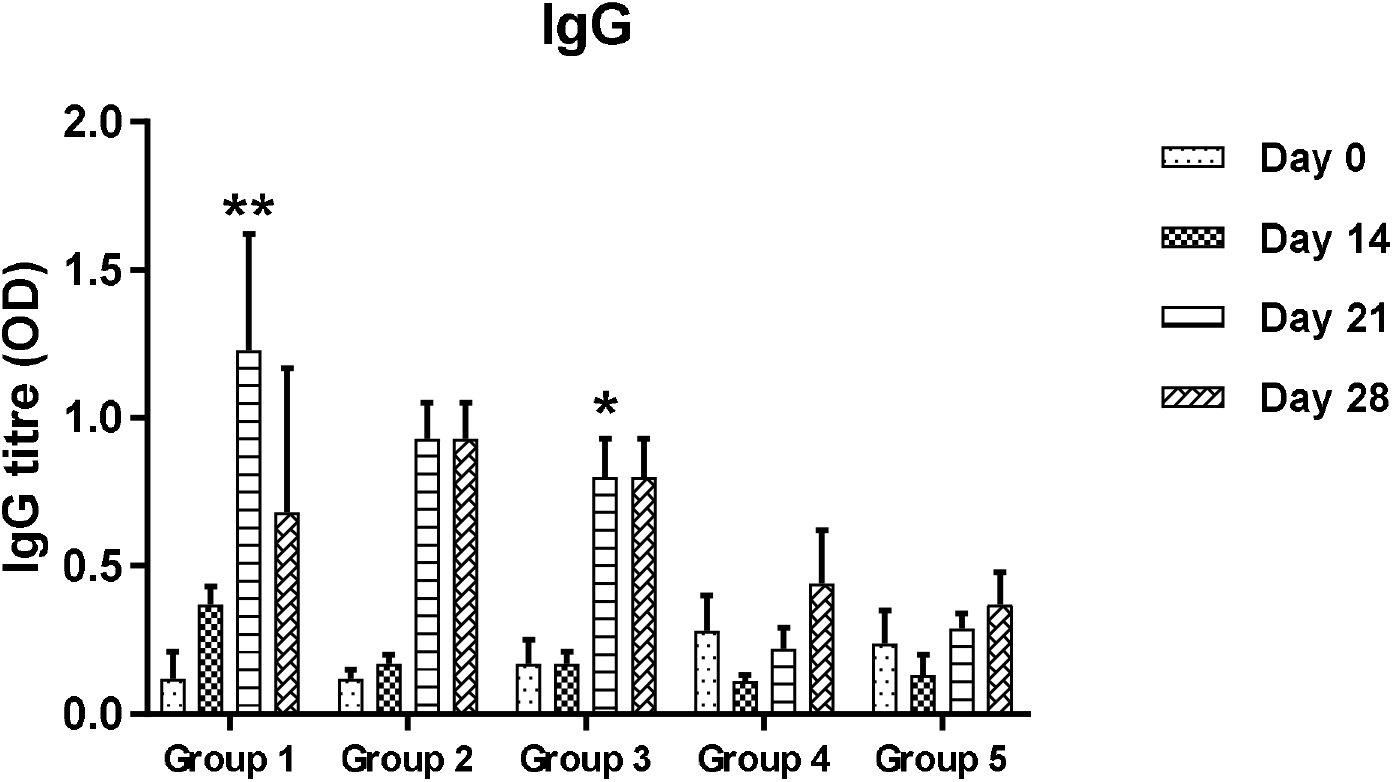
Effect of HBsAg-Chitin formulation on IgG titre of Mice Sera. (Results are expressed as mean + SD (n= 3). Values with **, * are significant at p < 0.05 (note; ** > *). (Group 1= [3 doses-HBsAg], Group 2= [2 doses-HBsAg]], Group 3 = [2 doses-HBsAg + Chitin], Group 4 = [2 doses-Chitin], Group 5 = [3 doses-Normal saline])

### 3.2. Effect of HBsAg-Chitin formulation on IgG1 titre of Mice Sera

**Figure 2** showed that, comparism between IgG_1_ titres of mice administered 3 doses of HBsAg and those administered HBsAg-Chitin formulation on days 14, 21 and 28, were statistically significant (p < 0.05). However, on day 21, IgG_1_ titres of mice administered HBsAg-Chitin formulation was significantly (p < 0.05) higher than those administered 2 doses of HBsAg. Moreover, On day 28, mice administered with 2 doses of HBsAg sustained a significantly (p < 0.05) higher IgG_1_ titre level when compared with mice administered 2 doses of HBsAg-Chitin formulation. IgG_1_ titres of mice administered 2 doses of HBsAg-Chitin formulation and those administered 2 doses of HBsAg increased significantly (p < 0.05) from days 0, through 14 to 21. However, IgG_1_ titres of mice administered 2 doses of HBsAg-Chitin formulation showed a significant (p < 0.05) decline on day 28.

**Figure 2:**
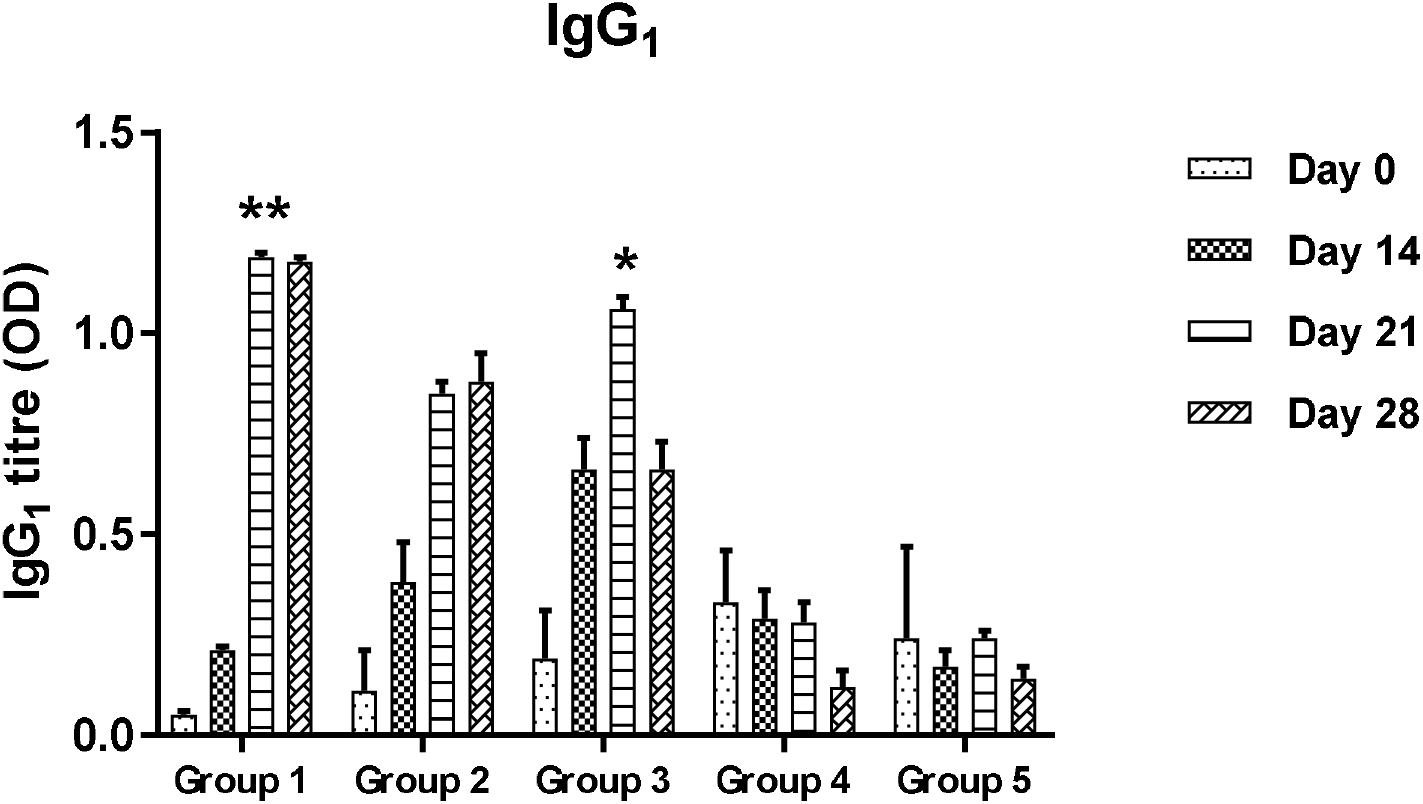
Effect of HBsAg-Chitin formulation on IgG1 titre of Mice Sera. (Results are expressed as mean + SD (n= 3). Values with **, * are significant atp < 0.05 (note; ** > *). (Group 1= [3 doses-HBsAg], Group 2= [2 doses-HBsAg]], Group 3 = [2 doses-HBsAg + Chitin], Group 4 = [2 doses-Chitin], Group 5 = [3 doses-Normal saline])

### 3.3. Effect of HBsAg-Chitin formulation on IgG2a Titre of Mice Sera

**Figure 3** showed that all treated mice groups did not elicit a significantly higher IgG2a titre when compared with the negative control (group 5).

**Figure 3:**
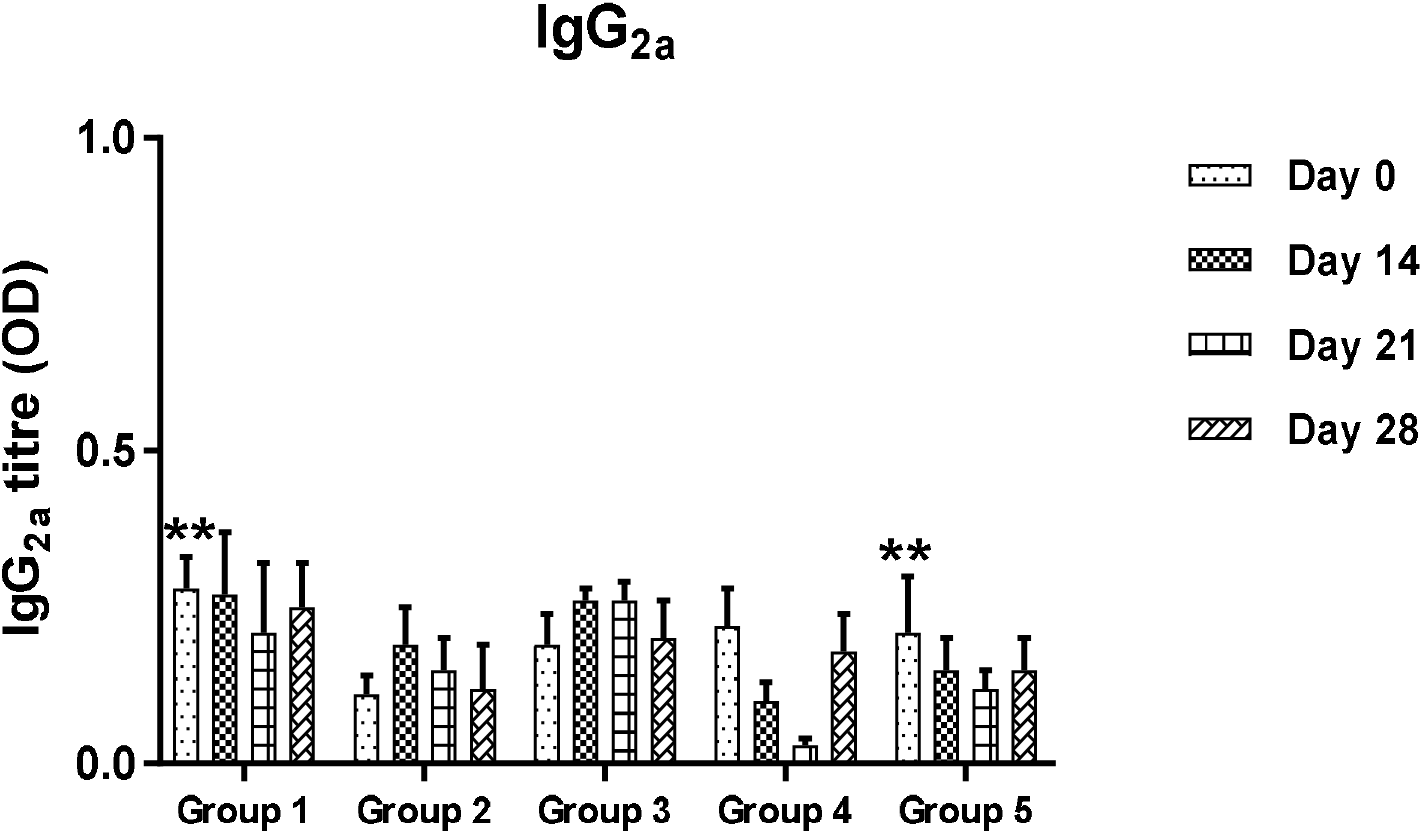
Effect HBsAg-Chitin formulation on IgG2a titre in Mice Sera. (Results are expressed as mean + SD (n= 3). Values with **, * are significant atp < 0.05 (note; ** > *). (Group 1= [3 doses-HBsAg], Group 2= [2 doses-HBsAg]], Group 3 = [2 doses-HBsAg + Chitin], Group 4 = [2 doses-Chitin], Group 5 = [3 doses-Normal saline])

### 3.4. Effect of HBsAg-Chitin formulation in Percentage Population of Lymphocytes in Mice

**Figure 4** illustrates apparently no significant (p > 0.05) increase or decrease in the percentage population of the lymphocytes across all groups. However, on day 28, there was a significant decline in percentage population of lymphocytes of all the treated groups relative to the negative control. Also, there was apparently a gradual decline in the mean percentage population of the lymphocytes in mice administered the HBsAg and HBsAg-Chitin formulation (see groups 1, 2, 3).

**Figure 4:**
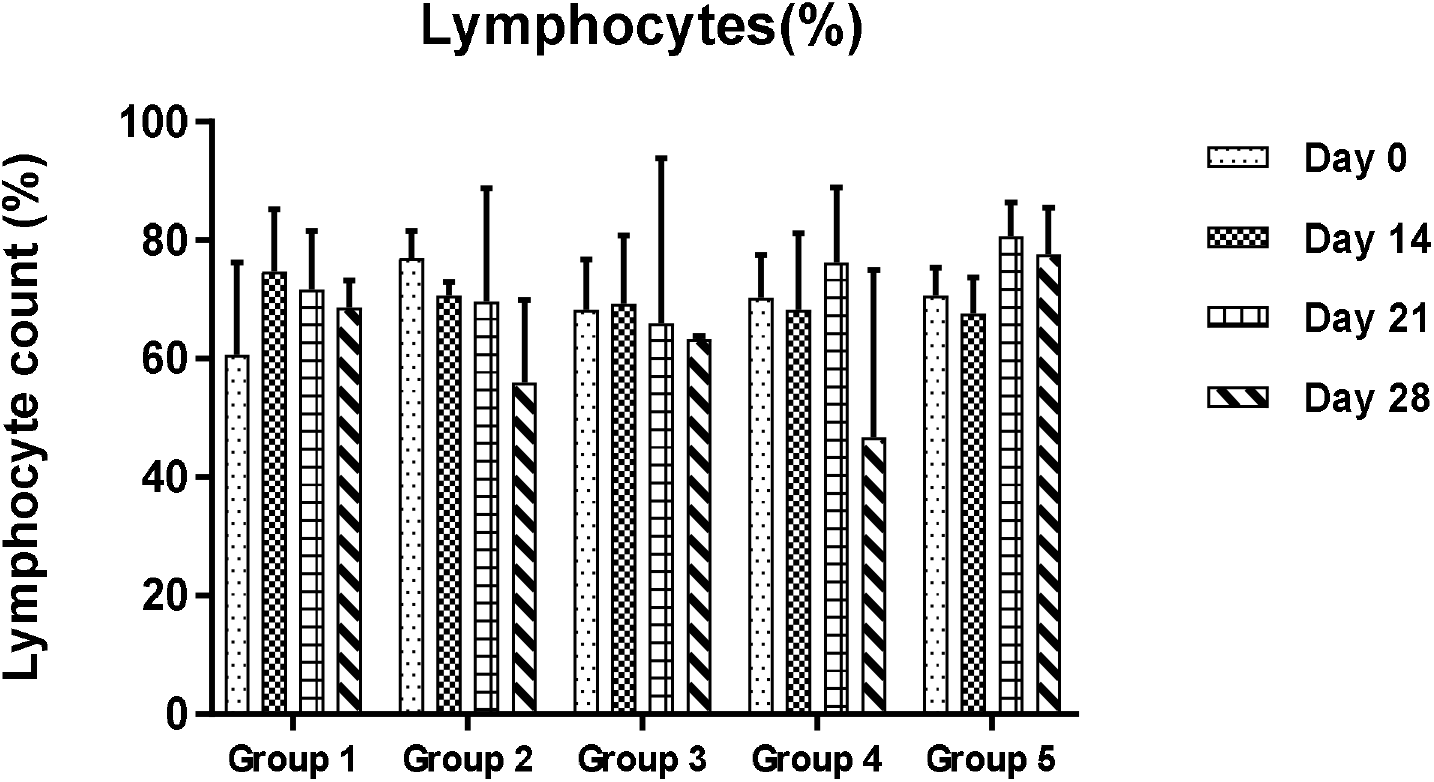
Effect of HBsAg-Chitin formulation in the percentage population of lymphocytes in mice sera. (Results are expressed as mean + SD (n= 3). Values with **, * are significant at p < 0.05 (note; ** > *). (Group 1= [3 doses-HBsAg], Group 2= [2 doses-HBsAg]], Group 3 = [2 doses-HBsAg + Chitin], Group 4 = [2 doses-Chitin], Group 5 = [3 doses-Normal saline]).

### 3.5. Effect of HBsAg-Chitin Formulation in the Percentage Population of Neutrophils in Mice

**Figure 5** showed that on days 21 and 28, the percentage population of neutrophils in the treated groups were significantly (p < 0.05) higher than that of the negative control (group 5). Apparently, the mean percentage population of the neutrophils increased gradually in mice administered the HBsAg and HBsAg-Chitin formulation after the first challenge (see groups 1, 2 3).

**Figure 5:**
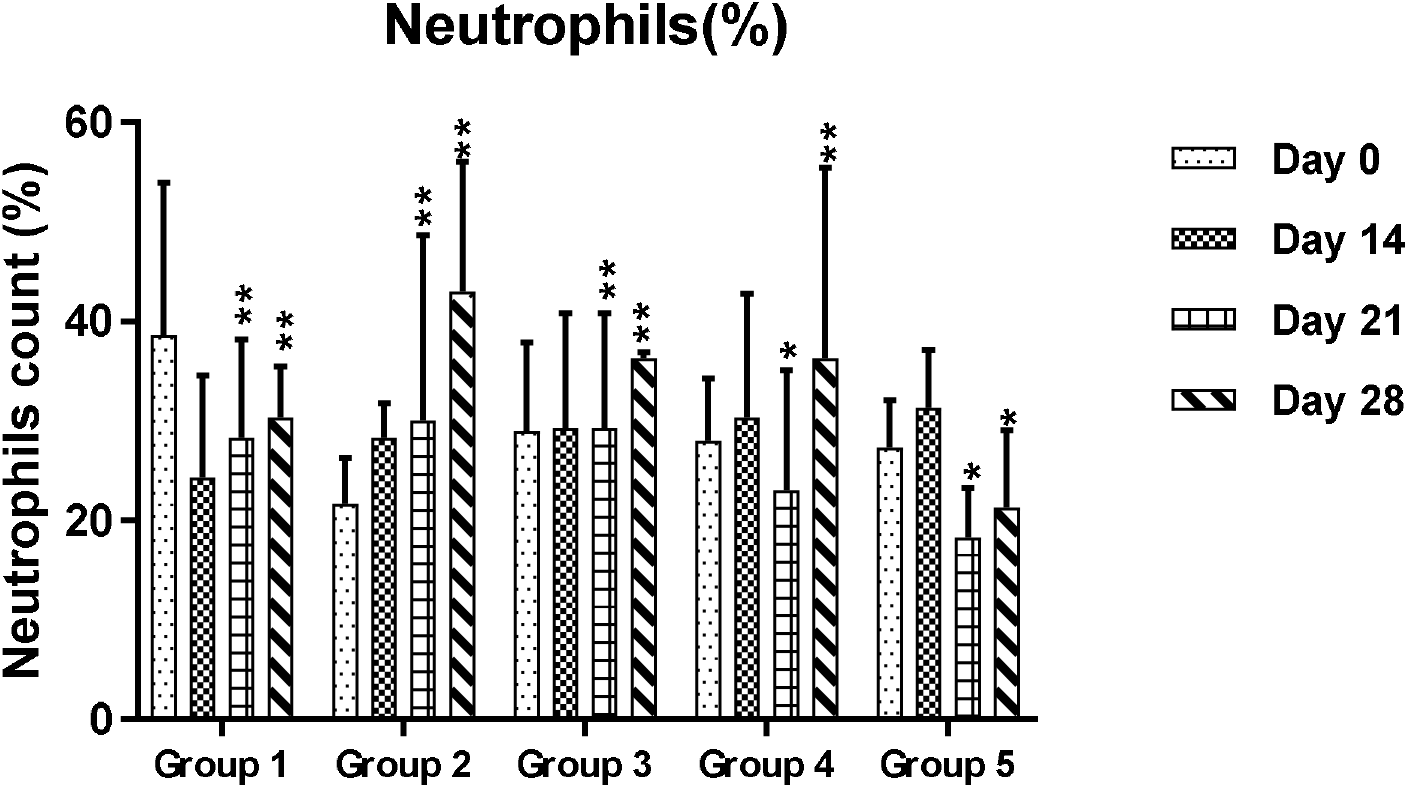
Effect of HBsAg-Chitin formulation in the percentage population of neutrophils in mice sera. (Results are expressed as mean + SD (n= 3). Values with **, * are significant at p < 0.05 (note; ** > *). (Group 1= [3 doses-HBsAg], Group 2= [2 doses-HBsAg]], Group 3 = [2 doses-HBsAg + Chitin], Group 4 = [2 doses-Chitin], Group 5 = [3 doses-Normal saline]).

### 3.6. Effect of HBsAg-Chitin formulation in Percentage Population of Monocytes in Mice

**Figure 6** showed that the percentage populations of monocytes of all treated groups declined significantly (p < 0.05) from day 14 to day 21 relative to the negative control (group 5). The mean percentage population of monocytes in mice administered with 2 doses of HBsAg-Chitin formulation (group 3) and chitin alone (group 4) were apparently much more down-regulated when compared with the positive control groups (groups 1 and 2).

**Figure 6:**
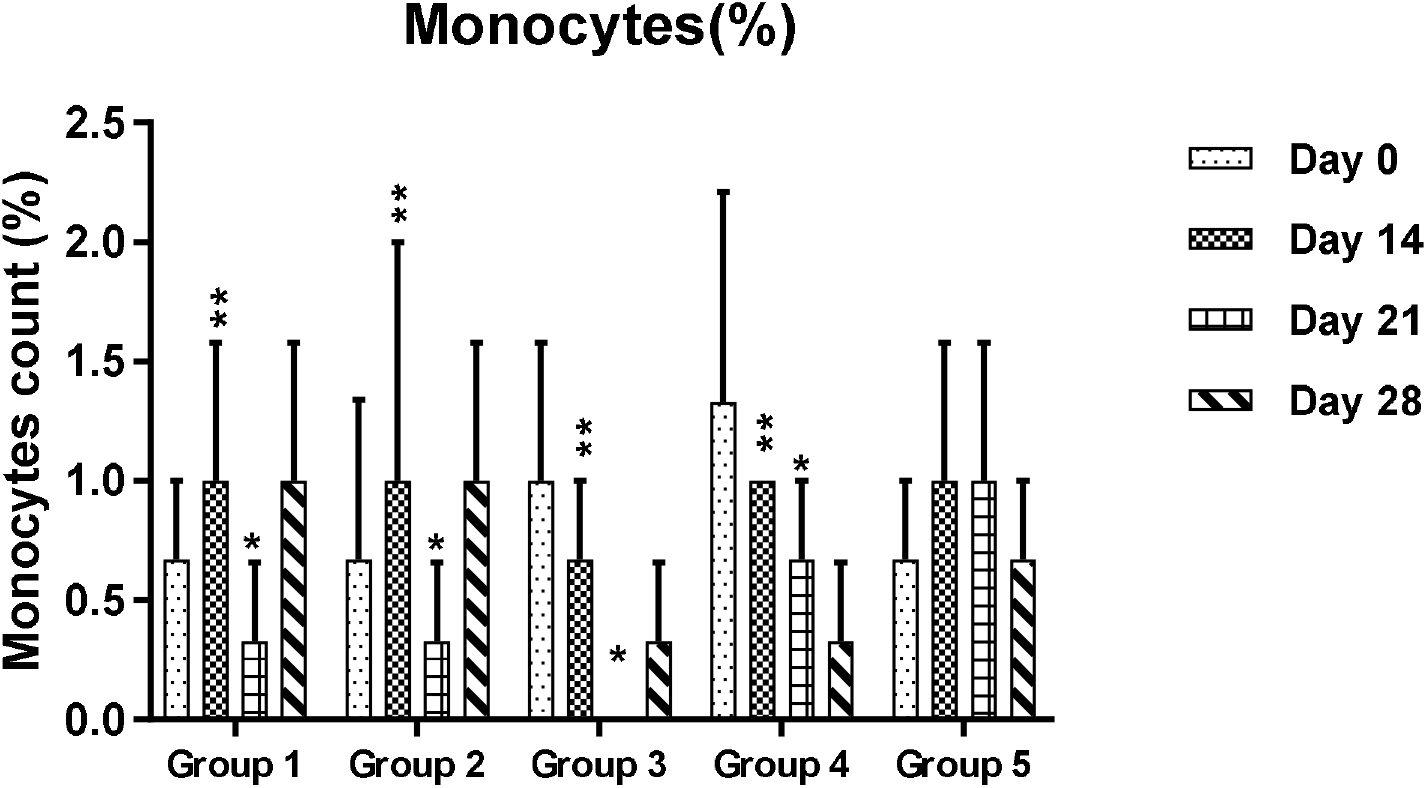
Effect of HBV-chitin combination in the percentage population of monocytes in mice sera. (Results are expressed as mean + SD (n= 3). Values with **, * are significant at p < 0.05 (note; ** > *). (Group 1= [3 doses-HBsAg], Group 2= [2 doses-HBsAg]], Group 3 = [2 doses-HBsAg + Chitin], Group 4 = [2 doses-Chitin], Group 5 = [3 doses-Normal saline]).

### 3.7. Effect of HBsAg-Chitin formulation in the Percentage Population of Eosinophils in Mice

**Figure 7** showed that, percentage population of eosinophils in mice administered 2 doses of HBsAg-Chitin formulation (group 3) was statistically (p < 0.05) different from those of all the other groups on day 0 through 14 to 21, though a decline in their population was observed on day 14. However, the mean percentage population of eosinophils declined significantly (p < 0.05) in all groups except on day 28 when there was a gradual increase in the eosinophil population in mice administered 2 doses of HBsAg-Chitin formulation (group 3) and the negative control (group 5)

**Figure 7:**
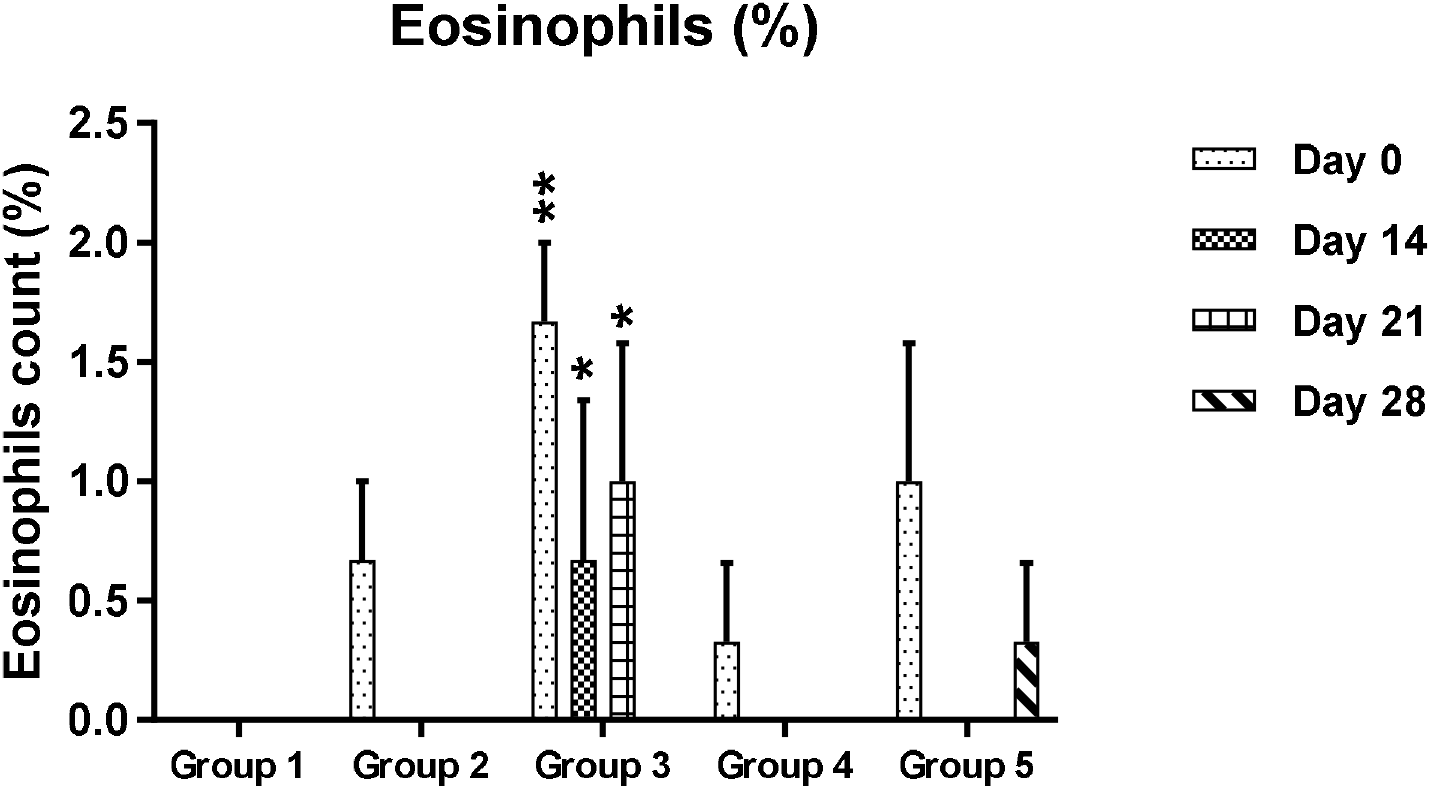
Effect of HBV-chitin in the percentage population of eosinophils in mice sera. (Results are expressed as mean + SD (n= 3). Values with **, * are significant at p < 0.05 (note; ** > *). (Group 1= [3 doses-HBsAg], Group 2= [2 doses-HBsAg]], Group 3 = [2 doses-HBsAg + Chitin], Group 4 = [2 doses-Chitin], Group 5 = [3 doses-Normal saline])

### 3.8. Effect of HBsAg-Chitin formulation on the Activities of Liver Marker Enzymes of Mice

**Table 1** illustrated that activities of alanine and aspartate aminotransferases (ALT, AST) of all experimental groups showed no statistically significant (p > 0.05) difference.

**Table 1:**
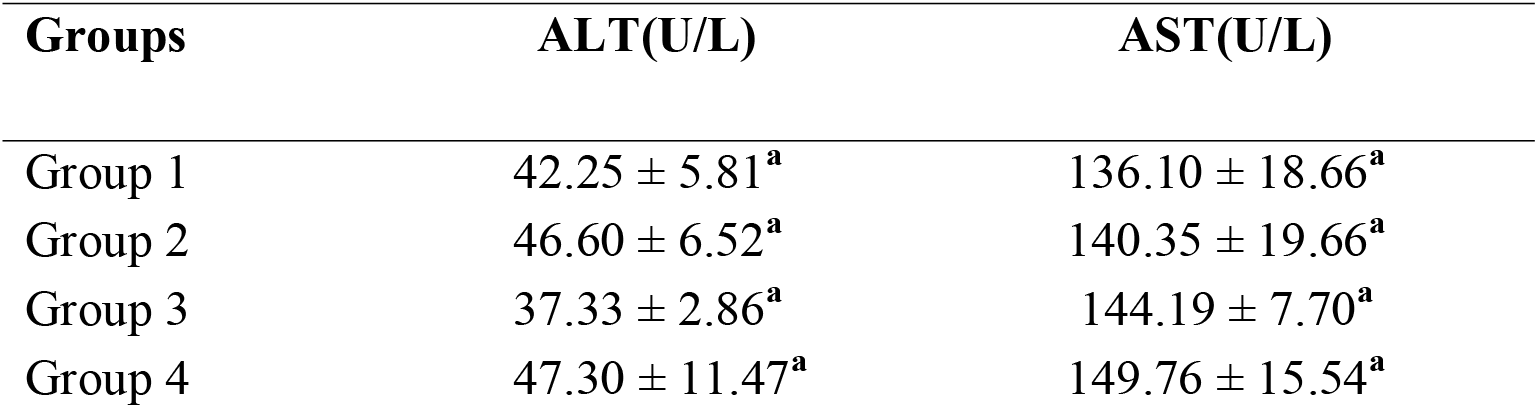
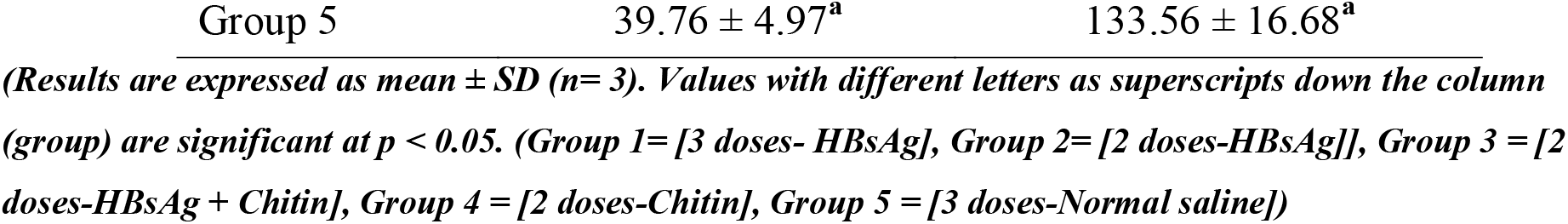
Effects of HBV-chitin on the activities of Liver Marker Enzymes of mice seras.

### 3.9. Effect of HBsAg-Chitin formulation in the Liver Weight of Mice

**Table 2** demonstrates that liver weight of mice that received 2 doses of HBsAg-Chitin formulation (group 3) was remarkably (p < 0.05) higher than that of mice that received 3 doses of HBsAg (group 1). However, the liver weight of mice administered with 2 doses of HBV-chitin combination (group 3) was not significantly (p > 0.05) higher when compared to the negative control group.

**Table 2:** Effect of HBV-chitin Combination on the Liver weight of Mice.

| Groups | Liver weight |
| --- | --- |
| Group 1 | $0.85 \pm 0.06^a$ |
| Group 2 | $1.07 \pm 0.17^{ab}$ |
| Group 3 | $1.33 \pm 0.20^c$ |
| Group 4 | $1.29 \pm 0.14^{bc}$ |
| Group 5 | $1.29 \pm 0.25^{bc}$ |

*Results are expressed as mean ± SD (n= 3). Values with different letters as superscripts ( $a < ab < bc < c$ ) down the column (group) are significant at $p < 0.05$ . (Group 1= [3 doses- HBsAg], Group 2= [2 doses-HBsAg], Group 3 = [2 doses-HBsAg + Chitin], Group 4 = [2 doses-Chitin], Group 5 = [3 doses-Normal saline])*

### 3.10. Mice body weight trend throughout the immunization schedules

**Fig. 8** shows relatively progressive increase in the mice body weights throughout the study period.

**Figure 8:**
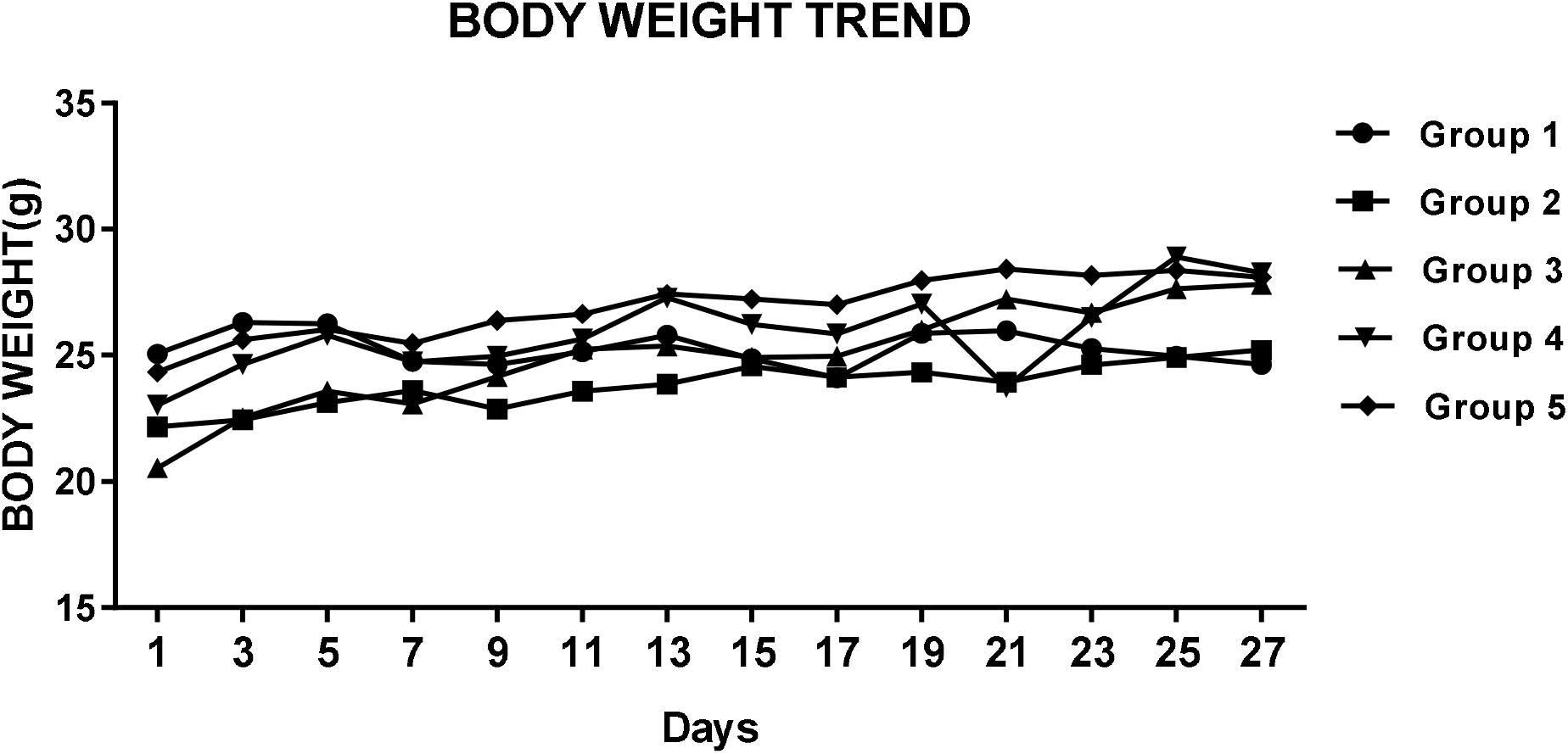
Mice body weight trend throughout the immunization schedules. (Group 1= [3 doses-HBsAg], Group 2= [2 doses-HBsAg]], Group 3 = [2 doses-HBsAg + Chitin], Group 4 = [2 doses-Chitin], Group 5 = [3 doses-Normal saline])

## 4. DISCUSSION

The humoral immune response was evaluated by measuring the titre levels of IgG, and its subclasses IgG1 and IgG2a in mice blood sera. The IgG class of immunoglobulin is generally produced in-vivo by memory B-cells after secondary exposure of the adaptive immune system to a recurring pathogen^19)^. Secondary exposure to an antigen is accompanied with a class switching from a previously circulating poorly avid immunolglobulin, IgM, to the IgG class that is more avid and binds with a much higher affinity and specificity to the antigen that elicited its production^20)^. In most vaccine clinical or preclinical studies, a highly elevated level of IgG class of immunoglobulins in blood serum correlates strong immunity to an anticipated infectious microorganism ^21)^. Strong neonatal immunity also gains credence to high IgG titre in fetal blood as the monomeric structure of IgG class of immunoglobulins permits its ability to permeate the placenta to protect the neonates ^19)^.

Previous claims about the adjuvant potentials accrued to chitin may be inconsistent as the results obtained from this study suggest the contrary^9,24,25)^. Our results reveal that HBsAg-Chitin formulation administered intramuscularly in a 2 dose schedule elicited a significantly (p < 0.05) lower IgG titre when compared with the HBsAg administered in a 3 dose schedule (figure 1). Conventionally, recombinant hepatitis-b vaccine is administered to subjects in a 3-dose schedule to achieve optimum immunity though it is not eternal ^6)^. An adjuvant that could help circumvent administration of the repeated booster doses and induce a more heightened and lasting immunity to pathogenic infection while maintaining the health safety status of a subject is the crux of adjuvant research^5)^. Seven days after the end of the vaccination period of this study (day 28), IgG titre level elicited by the 2 doses of HBsAg-Chitin formulation was non-significantly (p > 0.05) lower than that elicited by 2 doses of HBsAg (fig. 1). This suggests alum-based adjuvant which is a constituent of the HBsAg used in this study, may possess more promising adjuvant activity than chitin. This finding may negates the claim that chitin has excellent immunostimulatory activity^7)^. Chitin may seem to have shown slight adjuvant effect as the IgG titre level of mice administered with 2 doses of HBsAg-Chitin formulation showed a progressive rise from days 14 through days 21 to 28 (fig. 1). However, it will not be accurate to correlate the antibody titre elevation to the presence of chitin, since the observation was only within the group receiving the HBsAg-Chitin formulation. Considering the difference between the IgG titres of those receiving the HBsAg alone and the HBsAg-Chitin formulation, it is imaginable the presence of chitin was limiting the immunogenic potential of the recombinant hepatitis B vaccine. Possibly the presence of chitin altered the molecular patterns that are instrumental for pathogen recognition by the antigen presenting cells. We suggest that chitin may have formed a matrix that masked some of the molecular patterns displayed by the vaccine and consequently down-regulated optimum signaling of the antigen presenting cells (APCs) for adequate antigen processing.

Furthermore, observation from the IgG1 and IgG2a titres showed similar trends with the IgG titre. In mice, T-helper 2 (Th-2) mediated immune response has been associated with the induction of IgG1 production, while the induction of the production of IgG2a, IgG2b, and IgG3 by memory B-cells was associated with Th2-mediated immune response ^22)^. The major cytokine markers, IL-4, IL-5, and IL-13, secreted by the Th2 cells, stimulate B cell differentiation and subsequent antibody production to aid rapid clearance of extracellular pathogens^23)^. Our findings demonstrated that 2 doses of HBsAg-Chitin formulation (group 3) elicited significantly (p < 0.05) lower IgG_1_ titre level relative to 3 doses of HBsAg (group 1) alone (Fig. 2). The HBsAg-Chitin formulation did not also elicit the higher IgG_1_ titre than 2 doses of the recombinant hepatitis-B vaccine; hence it did not elicit a stronger Th2-mediated immune response.

All the treated groups demonstrated no significantly (p > 0.05) higher IgG2a titre relative to the negative control (group 5). This suggests that HBsAg and HBsAg-Chitin formulation may not activate T-helper 1 (Th-1) mediated immune responses. Th1-cells typically secrete interleukin-2 (IL-2) and interferon-gamma (IFN-γ) which are instrumental cytokines for the activation of macrophages and other cellular immune effectors that eliminate pathogens from intracellular sites ^26)^. This remains one of the limitations of the recombinant hepatitis b vaccine regarding stimulating rapid clearance of the virus from the intracellular milieu of the hepatocytes.

Moreover, the inability to elicit a stronger secondary immune response could also be influenced by the route of administration. Previous studies reported that chitin demonstrated a promising adjuvant activity when administered via the intranasal route ^24)^. Enhancement of vaccine action by chitin via administration through the mucosal route has been proposed to be mediated by intranasal retention of the vaccine and allowance for paracellular transport of the vaccine via the endothelial junctions^12)^. The mechanisms of vaccine enhancement by chitin through mucosal administration were believed to be due to both retention of vaccine in the nasal passages via mucoadhesion and opening of endothelial cell junctions for paracellular transport of vaccine ^27)^. However, the scope of this study did not address this phenomenon. The intramuscular route of administration may not have been appropriate for the HBsAg-Chitin formulation; however we aimed to closely mimick the mode of administration of the recombinant hepatitis-B vaccine as obtained in humans for a valid correlation ^6)^.

The cellular response of the immune system was evaluated by determining the percentage white blood cell differentials in mice peripheral blood. Immunization with alum and other adjuvants usually induce the mobilization of the granulocytes such as the polymorphonuclear-neutrophils (PMNs), eosinophils and basophils; the mononuclear monocytes and their macrophage derivatives, including the dendritic cells at the site of injection in a time-dependent manner ^28)^. The variation of the percentage of the lymphocytes among all the treated groups were not statistically significant (p>0.05), relative to the negative control (group 5). In the mice groups which received HBsAg and HBsAg-chitin formulation, there was apparently a decline in the lymphocyte population as days came by. This decline was independent of the vaccination as all the positive control groups followed the same trend. Peripheral lymphocyte count may not be a perfect correlate to vaccine outcomes. Recent study mentioned the unpredictability of hepatitis-b vaccine outcomes by evaluating regulatory B-lymphocyte levels in serum ^29)^.

An apparent non-significant (p > 0.05) increase in the mean percentage population of the neutrophils in all the treated groups from day 14 to 28 indicates the mobilization of neutrophils against foreign agents that enter the body ^30)^. There seemed to be a competition between the neutrophils and lymphocytes suggesting that granulocyte colony-stimulating factor (GCSF) may have been secreted and brought about preferential recruitment of neutrophils over the lymphocytes as days came by ^31)^. The increase in the percentage population of neutrophils was associated with a corresponding decline in the lymphocyte levels. This may be related to concurrent decline of lymphocytes as antigens are gradually cleared and processed by the antigen presenting cells which are said to be down-regulated upon interaction with the neutrophils. Previous study had reported interference with antigen presentation due to competition between the neutrophils and dendritic cells, which in turn regulates the mobilization of lymphocytes ^32)^.

A significant (p > 0.05) decline in the percentage population of the monocytes within all the treated groups on day 21, further suggests that the immune response to the vaccine and the reconstituted vaccine was not Th1-biased. Macrophages which are differentiation derivatives of blood monocytes are activated in the presence of some cytokines (IL-2, IFN-γ) secreted by the Th1-cells ^33)^. Hence, intracellular clearance of the hepatitis b virus may not have been achieved.

Although the percentage population of the eosinophils in mice that received the HBsAg-Chitin formulation was significantly (p < 0.05) higher than other groups, this variation was independent of the treatment received by each group; since the baseline (Day 0) eosinophil percentage population of the mice that received the HBsAg-Chitin formulation was quite higher relative to all the other groups. According to Ó Connell et al. (2015), the highest percentage population of eosinophils in mice is approximately 7%, which in our results revealed that it did not deviate from the norm^34)^. Apparently, hypersensitive reaction associated with elevated levels of eosinophils in blood was not obtainable in all the mice groups, and consequently we affirm chitin’s biocompactibility and nontoxicity^7)^. Also basophils were not observed in all mice groups throughout the study, which further validates the claim of rare presence of basophils in mice blood ^34)^. Basophils and eosinophils generally initiate immune responses by secreting pro-inflammatory mediators in the presence of infectious agents.

Hepatic injury can be assessed by measuring levels of alanine and aspartate aminotransferases (ALT, AST), alkaline phosphatase (ALP) in blood serum ^35)^. Leakage of these hepatic intracellular enzymes into the blood circulation upsets liver homeostasis and serves as metabolic indicators of hepatitis or other forms of liver damage ^36)^. In humans, a patient with an AST:ALT ratio > 2:1 is indicative for liver metabolic disorder, and an AST:ALT ratio > 3:1 is strongly indicative for alcoholic liver disease ^37)^. Low level of ALT in the serum is associated with deficiency of pyridoxal phosphate, which was reported to be induced by lethal doses of alcohol. As shown in Table 1, ALT and AST levels were not significantly (p > 0.05) different across all treated groups, with apparently similar liver weight values (Table 2), which suggests that HBsAg-Chitin formulation may exert no hepatotoxic effect on the liver and was further supported by an overall body weight gain of the mice across all groups.

Moreover, considering the chitin’s size-dependent biological functions we suggest that formulating recombinant protein vaccine with chitin nanoparticles and lesser quantity than was used in this study may improve its adjuvanticity. Chitin apparently possesses chelating property which masked molecular patterns detectable by the antigen presenting cells and in turn could not initiate stronger signals to activate antibody production.

In conclusion, this present study demonstrated that chitin in combination with recombinant hepatitis b protein-alum vaccine apparently elicited a weaker secondary humoral immune response relative to HBsAg, though considered to be safe.

## References

1. Manzoor S., Saalim M., Imran M., Resham S. and Ashraf J. Hepatitis B virus therapy: What's the future holding for us? World J Gastroentero., 21, 12558–12575 (2015).

2. O'Hagan D. T. and Gregorio E. D. The path to a successful vaccine adjuvant. Drug Discov., 14, 541–551 (2009).

3. Pasquale A.D., Preiss S., Da-Silva F.T. and Garçon N. Vaccine adjuvants: from 1920 to 2015 and beyond. Vaccines, 3, 320–343 (2015).

4. O'Hagan D. T. and Rappuoli R. Novel approaches to vaccine delivery. Pharmacol. Res., 21, 1519–1530 (2004).

5. Lee S. and Nguyen M. T. Recent advances of vaccine adjuvants for infectious diseases. Immune Netw., 15(2), 51–57 (2015).

6. Leuridan E. and Damme P. V. Hepatitis B and the need for a booster dose. Vaccines, 53, 68–75 (2011).

7. Li X., Min M., Du N., Gu Y., Hode T., Naylor M., Chen D., Nordquist R. E., and Chen W. R. Chitin, chitosan, and glycated chitosan regulate immune responses: the novel adjuvants for cancer vaccine. Clin. Dev. Immunol., 2013, 1–8 (2013).

8. Jarmila V. and VavŕJková E. Chitosan derivatives with antimicrobial, antitumour and antioxidant activities-a review. Curr. Pharm. Des., 17 (32), 3596–3607 (2011).

9. Nishimura K., Nishimura S., and Nishi N. Immunological activity of chitin and its derivatives. Vaccine, 2 (1), 93–99 (1984).

10. Tokura, S., Tamura, H., and Azuma, I. Immunological aspects of chitin and chitin derivatives administered to animals. EXS, 87, 279–292 (2005).

11. Y. Shibata, L. A. Foster, W. J. Metzger et al., Alveolar macrophage priming by intravenous administration of chitin particles, polymers of N-acetyl-D-glucosamine, in mice. Infect. Immun., 65 (5), 1734–1741 (1997).

12. Shibata Y., Metzger J. W. and Myrvik Q. N. Chitin particle-induced cell-mediated immunity is inhibited by soluble mannan: mannose receptor-mediated phagocytosis initiates IL-12 production. J Immunol, 159 (5), 2462–2467 (1997).

13. Islam S. Z., Khan M. and Nowsad Alam A. K. M. Production of chitin and chitosan from shrimp shell wastes. J. Bangladesh Agril. Univ., 14 (2), 253–259 (2016).

14. National Research Council Institute for Laboratory Animal Research. Guide for the care and use of laboratory animals. National Academy Press, Washington DC (1996).

15. Ternette N., Tippler B., Überla K. and Grunwald T. Immunogenicity and efficacy of codon optimized DNA vaccines encoding the F-protein of respiratory syncytial virus. Vaccine, 25, 7271–7279 (2007).

16. Rodak B. F. and Carr J. H. Clinical Hematology Atlas. 3rd edition, Elsevier, Saunders, p. 296 (2016).

17. Thomas L. Alanine aminotransferase (ALT), Aspartate aminotransferase (AST). Clinical Laboratory Diagnostics. (Thomas L. ed) 1st edition, TH-Books Verlagsgesellschaft, Frankfurt, pp. 55–65 (1998).

18. Moss D. W. and Henderson A. R. Clinical enzymology. Tietz Textbook of Clinical Chemistry. (Burtis C. A. and Ashwood E. R. eds.) 3rd edition, W.B. Saunders Company, Philadelphia, pp. 617–721 (1999).

19. Willey J. M., Sherwood L. M. and Woolverton C. J. Prescott's microbiology, 9th edition. McGraw-Hill Education, New York, (2014).

20. Burns R. Immunochemical techniques. Principles and techniques of biochemistry and molecular biology. (Wilson K. and Walker J. eds.) 7th edition, Cambridge University Press, Cambridge, pp. 263–299 (2010).

21. Alvarado-Mora M. V., Fernandez M. F., Gomes-Gouvêa M. S., de Azevedo Neto R. S., Carrilho F. J. and Pinho J. R.. Hepatitis B (HBV), hepatitis C (HCV) and hepatitis delta (HDV) viruses in the Colombian population--how is the epidemiological situation? PLoS One, 6(4), e18888 (2011).

22. Germann T, Bongartz M, Dlugonska H, Hess H, Schmitt E, Kolbe L, Kolsch E, Podlaski FJ, Gately MK, Rude E. Interleukin-12 profoundly up-regulates the synthesis of antigen-specific complement-fixing IgG2a, IgG2b and IgG3 antibody subclasses in vivo. Eur. J. Immunol., 25(3), 823–829 (1995).

23. Andris F., Denanglaire S., Anciaux M., Hercor M., Hussein H. and Leo O. The transcription factor c-maf promotes the differentiation of follicular helper T cells. Front. Immunol., 8,480 (2017).

24. Read R. C., Naylor S. C., Potter C. W. et al. Effective nasal influenza vaccine delivery using chitosan. Vaccine, 23 (35), 4367–4374, (2005).

25. McNeela E. A., Jabbal-Gill I., Illum L. et al. Intranasal immunization with genetically detoxified diphtheria toxin induces T cell responses in humans: enhancement of Th2 responses and toxin-neutralizing antibodies by formulation with chitosan. Vaccine, 22 (8), 909–914 (2004).

26. Biedermann T., Zimmermann S., Himmelrich H. IL-4 instructs TH1 responses and resistance to Leishmania major in susceptible BALB/c mice. Nat. Immunol., 2, 1054–1060 (2010).

27. Amorij J. P., Hinrich W. L. J., Frijlink H. W., Wilschut J. C. and Huckriede A. Needle-free influenza vaccine. Lancet Infect. Dis., 10, 699–711 (2010).

28. Lu F. and Hogenesch H. Kinetics of the inflammatory response following intramuscular injection of aluminium adjuvant. Vaccine, 31 (37), 3979–3986 (2013).

29. Maria B., Karen L. D., Martin T., Kumar V., Lan W., Narelle S. et al. Levels of regulatory B cells do not predict serological responses to hepatitis B vaccine. Hum. Vaccin. Immunother., 14, 1483–1488 (2018).

30. Ghimire T. R., Benson R. A., Garside P. and Brewer J. M. Alum increases antigen uptake, reduces antigen degradation and sustains antigen presentation by dendritic cells in vitro. Immunol. Lett., 147(2), 55–62 (2015).

31. Melve G. K., Ersvaer E., Eide G. E., Kristoffersen E. K. and Bruserud Ø.. Peripheral blood stem cell mobilization in helthy donors hy granulocute colony-stimulating factor causes preferential mobilization of lymphocyte subsets. Front Immunol, 9:845 (2018).

32. Yang C. W., Strong B. S., Miller M. U., Unanue E. R. Neutrophils influence the level of antigen presentation during the immune response to protein antigens in adjuvants. J Immunol., 185 (5), 2927–2934 (2010).

33. Leopold W. C. and Wormley F. Classical alternative macrophage activation: the ying and yang in host defense against pulmonary fungal infections. Mucosal Immunol, 7, 1023–1035 (2014).

34. Ó Connell K. E., Mikkola A. M., Stepanek A. M., Vernet A., Hall C. D, Sun C. C., Yildirim E., Staropoli J. F., Lee J. T., Brown D. E. Practical murine hematopathology: a comparative review and implications for research. Comparative Med, 65, 96–113 (2015).

35. Pincus M. R. and Schaffer J. A. Assessment of liver function. Clinical Diagnosis and Management by Laboratory Methods (Henry J. B. ed.). 20th edition, W.B. Saunders Company, pp. 253–267 (2013).

36. Nyblom, H., Bjornsson, E., Simren, M., Aldenborg, F., Almer, S., Olsson, R. The AST/ALT ratio as an indicator of cirrhosis in patients and establish a prognosis. Postgrad. Med. J., 107 (2), 113–114 (2012).

37. Ni. H., Soe H.H.K. and Htet A. Determinants of abnormal liver function tests in diabetes patients in Myanmar. Int. J. Diabetes Res., 1 (3), 34–41 (2012).

38. Goffin E, Horsmans Y, Cornu C, Geubel A, Pirson Y. Acute hepatitis B infection after vaccination. Lancet, 345 (8944), 261–263 (1995).

